# Asparagine availability is an essential limiting factor for poxvirus protein synthesis

**DOI:** 10.1101/427203

**Authors:** Anil Pant, Shuai Cao, Zhilong Yang

## Abstract

Virus actively interfaces with host metabolism because viral replication relies on host cells to provide nutrients and energy. For efficient viral replication in culture, vaccinia virus (VACV; the prototype poxvirus) prefers glutamine to glucose, to the extent that in glutamine-free medium, VACV replication is inefficient. Remarkably, VACV replication can be fully rescued from glutamine depletion by asparagine supplementation. By global metabolic profiling, genetic and chemical intervening of asparagine supply, we provide evidence demonstrating that the requirement of asparagine for efficient viral replication accounts for VACV’s preference of glutamine to glucose, rather than because glutamine is superior to glucose in feeding the tricarboxylic acid (TCA) cycle. Further, we show that asparagine availability is a critical factor for efficient viral protein synthesis. Our study highlights that the asparagine metabolism, whose regulation has been evolutionarily tailored in mammalian cells, presents a critical barrier to poxvirus replication, suggesting new directions of anti-viral strategy development.

---

Viruses do not have metabolisms; so viral replication relies on the supply and availability of host nutrients and energy. Not surprisingly, metabolism is a key interface of virus-host interactions, with many viral infections characterized by requirements for particular metabolites (e.g., glutamine, glucose or fatty acids) for optimal replication. Conceivably, many viruses induce alterations in metabolic pathways such as glycolysis, synthesis of fatty acids and nucleotides, and energy metabolism, and these changes shape the outcomes of viral infections ^1–4^.

Vaccinia virus (VACV) is the prototype poxvirus, with a large double-stranded DNA genome that encodes more than 200 annotated genes ^5,6^. Many poxviruses cause fatal diseases, including smallpox, one of the most devastating infectious diseases in human history. Although eradicated in nature, smallpox is still a valid national-security concern through potential unregistered stocks or *de novo* synthesis of live variola virus, the causative agent of smallpox ^7–9^. Poxviruses not only cause deadly epidemics, however, because poxviruses are practically useful as oncolytic agents for cancer treatments, vectors for vaccine development, and recombinant protein production ^10–13^. For efficient VACV replication in cell culture, VACV prefers glutamine to glucose, i.e. depletion of glutamine, but not glucose, from culture medium significantly decreases VACV production ^14,15^. In line with this finding, VACV infection upregulates glutamine metabolism ^16^. Nevertheless, why VACV prefers glutamine to glucose for replication remains unknown.

Glutamine is a non-essential amino acid that is abundantly utilized by mammalian cells beyond its role as a protein building block ^17^. Glutamine feeds the tricarboxylic acid cycle (TCA cycle; **Fig. 1A**) through glutamate and alpha-ketoglutarate (α-KG), in a process known as anaplerosis ^18–20^. Glutamine also acts as a biosynthetic precursor for many molecules, including amino acids, nucleotides, and fatty acids ^21,22^. Although several non-essential amino acids require intermediates of glutamine metabolism for *de novo* biosynthesis, only asparagine biosynthesis exclusively depends on glutamine because the amination of the synthesis reaction requires glutamine ^23,24^. The biosynthesis of asparagine using glutamine is catalyzed by the enzyme asparagine synthetase (ASNS) ^25,26^.

**Figure 1.**
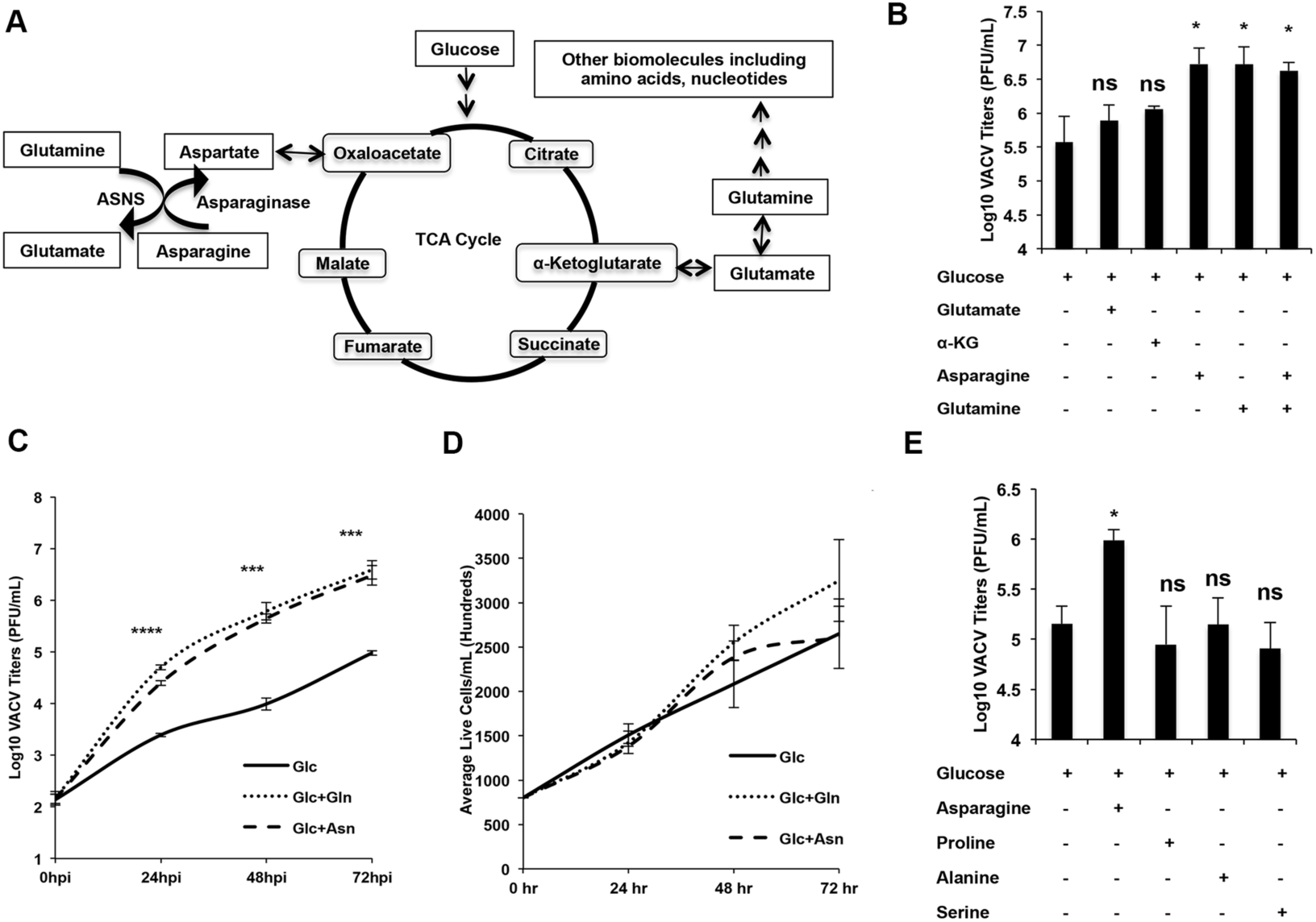
Asparagine fully rescues VACV replication from glutamine depletion. **(A)** Schematic of the role of glutamine in the TCA cycle and biomolecule synthesis. Note asparagine is the only amino acid that exclusively requires glutamine for its biosynthesis. **(B)** Asparagine fully rescues VACV replication from glutamine depletion, while α-KG and glutamate do not. HFFs were infected with VACV at an MOI of 2 in media containing 1 g/L glucose, 2 mM glutamine, 2 mM asparagine, 7 mM α-KG, or 5 mM glutamate as indicated. VACV titers were measured by a plaque assay at 24 hpi. **(C)** Asparagine rescues VACV growth kinetics from glutamine depletion. HFFs were infected with VACV at an MOI of 0.001 in medium containing the indicated nutrients. VACV titers were measured by a plaque assay at the indicated time, post infection. **(D)** HFF proliferation is not affected in different media. Equal numbers of HFFs were seeded in the indicated media. The cell numbers were counted over a period of 72 h using a hemocytometer. **(E)** Proline, alanine and serine cannot rescue VACV replication from glutamine depletion. Experiments were carried out similar to (B) with 5 mM proline, 1 mM alanine or 1 mM serine used. Error bars represent the standard deviation between at least three biological replicates. ns=p > 0.05, * = p ≤ 0.05, *** = p ≤0.001 and **** = p ≤0.0001.

A new and growing body of work suggests asparagine is more than a polypeptide subunit but is also essential in coordinating overall protein synthesis, cellular responses to amino acid homeostasis, and metabolic availability during biological processes and disease development. For example, asparagine acts as a metabolic regulator of TCA cycle intermediates and the cellular supply of nitrogen (which supports the synthesis of non-essential amino acids); and for cancer cells, asparagine bioavailability is essential for survival, proliferation and tumor development ^23,24,27,28^. Asparagine is also important for supporting kaposi’s sarcoma-associated herpes virus (KSHV) transformed cancer cell proliferation during glutamine depletion ^29^. However, the role of asparagine and its availability in virus replication has not been explored.

In the current study, we show that asparagine is a limiting metabolite for VACV replication through its critical role in VACV post-replicative mRNA translation. In contrast to the generic paradigm that glutamine is a superior to glucose in fueling the TCA, we show that the preference for glutamine reflects the asparagine requirement during replication. Indeed, interfering with asparagine metabolism severely impaired VACV replication, highlighting the importance of asparagine availability during the VACV lifecycle. Our findings demonstrate the essential role of asparagine in VACV replication. This critical host-dependent barrier to VACV replication might not only spur development of new, host-oriented antiviral therapies but also improve development of poxviruses as therapeutic tools.

## Results

### Asparagine fully rescues VACV replication from glutamine depletion

To test why VACV prefers glutamine to glucose for efficient replication, we examined whether α-KG and glutamate—the products of glutaminolysis that feed the TCA cycle (**Fig. 1A**)—could rescue VACV replication from glutamine depletion. Measuring virus titers showed that, in the absence of glutamine, α-KG and glutamate only partially or not at all rescued VACV replication, in agreement with an earlier study (**Fig. 1B**) ^14^. This indicates that while anaplerosis of the TCA cycle is important, this function of glutamine is not responsible for its superiority to glucose in promoting VACV replication.

Notably, VACV replication was fully rescued from glutamine depletion when asparagine was added to the medium (**Fig. 1B**). In contrast, when the medium contained glutamine, adding asparagine did not boost viral titers, suggesting growth could be equally rescued by either glutamine or asparagine.

Normal VACV replication kinetics in glutamine-free medium was also consistently rescued by asparagine (over a 72-h replication period, when the initial VACV multiplicity of infection (MOI) was 0.001; **Fig. 1C**). Accordingly, in VACV-infected cells grown in the absence of glutamine, asparagine rescued GFP expression from the VACV late promoter (**Fig. S1**). In medium containing glucose only, the 72-h proliferation rate of HFFs differed little from cells grown in glucose plus glutamine, or glucose plus asparagine (**Fig. 1D**), suggesting the difference in VACV titers is not due to cytotoxicity or altered HFF proliferation. Other non-essential amino acids that can be synthesized from glutamine, but were not present in the cell culture medium (e.g., proline, alanine, and serine), did not rescue VACV replication from glutamine-deficiency (**Fig. 1E**). Taken together, our results demonstrate that asparagine can fully rescue VACV replication from glutamine depletion.

### Asparagine does not enhance TCA cycle activities during VACV infection

Under glutamine-free conditions, asparagine might hypothetically rescue VACV replication by enhancing anaplerosis of the TCA cycle. To test this idea, VACV-infected HFFs were profiled for metabolic activities in three different conditions: glucose, glucose plus glutamine, and glucose plus asparagine. Here, compared to glucose-only conditions, glucose plus glutamine significantly enhanced the concentrations of several TCA cycle intermediates (α-KG, malate, fumarate and succinate; **Fig. 2A**), while addition of asparagine did not (**Fig. 2A**). Even in the absence of glutamine, in fact, glucose was sufficient to maintain the levels of oxidative-phosphorylation intermediates required for ATP production (**Fig. 2B**). These results show rescue of VACV replication in glutamine-deficient medium is not driven by enhancement of the TCA cycle, and that glucose can support enough TCA cycle activities for VACV infection. This data is supported by the results showing that by inhibiting glutaminase activity (with the specific inhibitor BPTES) VACV titers decreased by only 2-fold in the presence of glucose, but it decreased by 12-fold in the absence of glucose (glutamine was present in both conditions; **Fig. 2C**). Together these results indicate that, in glutamine-depleted conditions, asparagine-mediated rescue of VACV replication is not due to enhanced TCA cycle activity.

**Figure 2.**
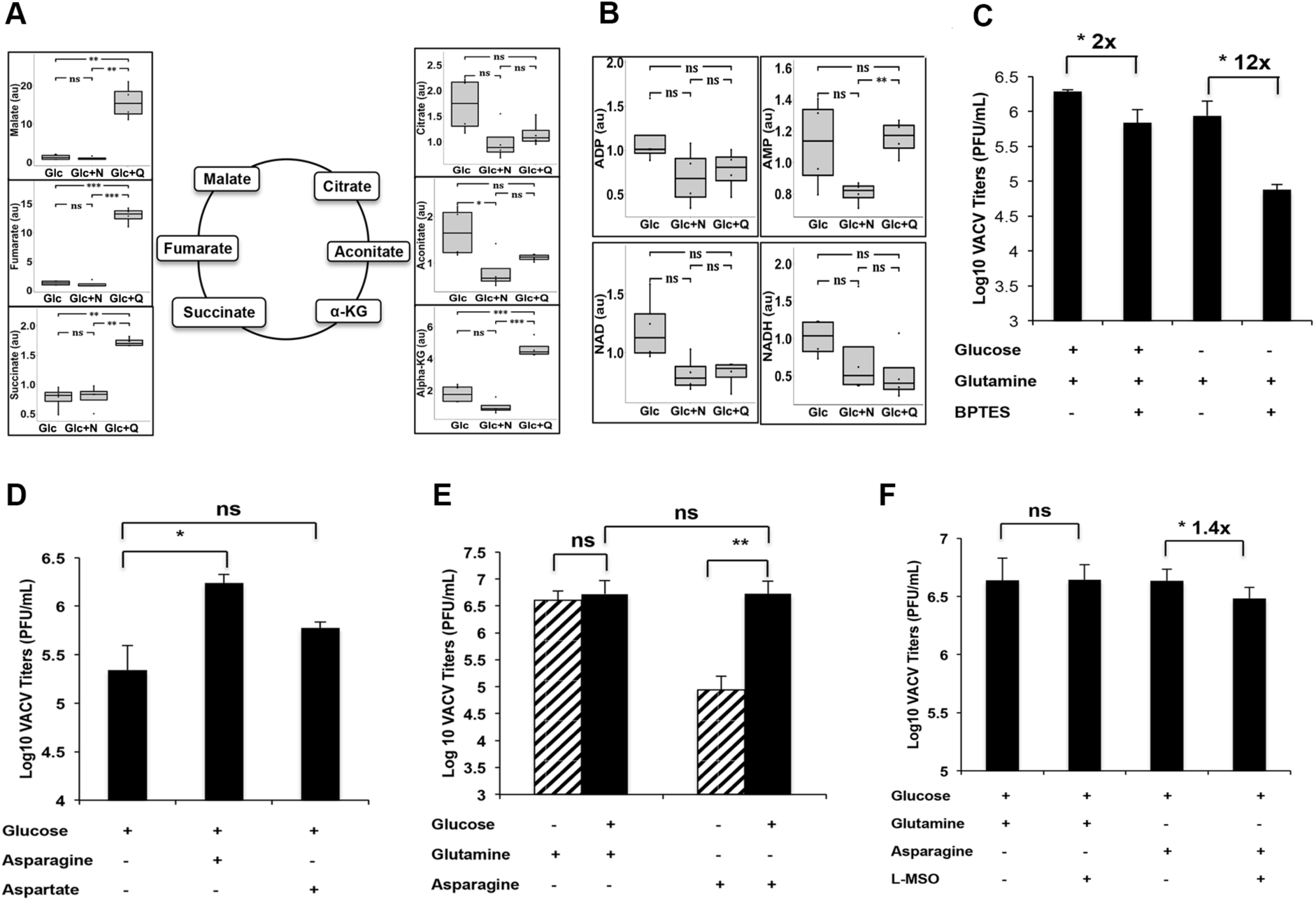
Asparagine supplementation does not enhance TCA cycle activities under glutamine-depleted conditions during VACV infection. **(A)** Asparagine addition does not recapitulate glutamine’s effect on TCA cycle activities. Levels of TCA cycle intermediates in HFFs infected with VACV (MOI of 3) for 8 h, in media containing glucose (Glc), glucose plus asparagine (Glc+N), or glucose plus glutamine (Glc+Q), determined by a global metabolic profiling. **(B)** Levels of oxidative phosphorylation intermediates in VACV-infected HFFs are not significantly different (from the same metabolic profiling as described in A). **(C)** Inhibiting glutaminase activity more severely affected VACV replication in the absence of glucose. HFFs infected with VACV, at an MOI of 2 for 24 h, in medium containing 10 μΜ BPTES or DMSO (control). The numbers indicate the fold change in VACV titer compared to DMSO treatment. **(D)** Aspartate is not as efficient as asparagine in supporting VACV replication. VACV titers in HFFs infected with VACV, at an MOI of 2 for 24 h, in indicated medium were measured by a plaque assay. **(E)** Rescue of VACV replication from glutamine depletion requires the presence of glucose. HFFs were infected with VACV, at an MOI of 2 for 24 h, in indicated media. VACV titer was measured by a plaque assay. **(F)** Inhibition of glutamine synthetase only mildly affected VACV replication. HFFs infected with VACV, at an MOI of 2 for 24 h, in indicated medium containing 2 mM L-Methionine Sulfoximine (MSO) or DMSO. VACV titers were determined by a plaque assay. Error bars represent the standard deviation between four biological replicates (Fig 2a, 2b) and at least three biological replicates (Fig 2c-2f). ns=p > 0.05, *= p ≤ 0.05, **= p ≤0.01, ***= p ≤0.001. The numbers above the bars represent fold change.

Our interpretation is also supported by the fact that, in VACV-infected cells, the TCA cycle is not directly fed by asparagine converting to aspartate: adding aspartate to glutamine-deficient medium rescued only low levels of VACV replication (**Fig. 2D**); adding asparagine did not elevate aspartate concentration (**Fig. 3A**), which is consistent with the fact that in mammalian cells asparaginase does not actively convert asparagine to aspartate ^28^. Notably, glutamine, itself, could support VACV replication, even in the absence of glucose, while asparagine-mediated rescue of VACV replication, from glutamine depleted conditions, required glucose in the medium (**Fig. 2E**).

**Figure 3.**
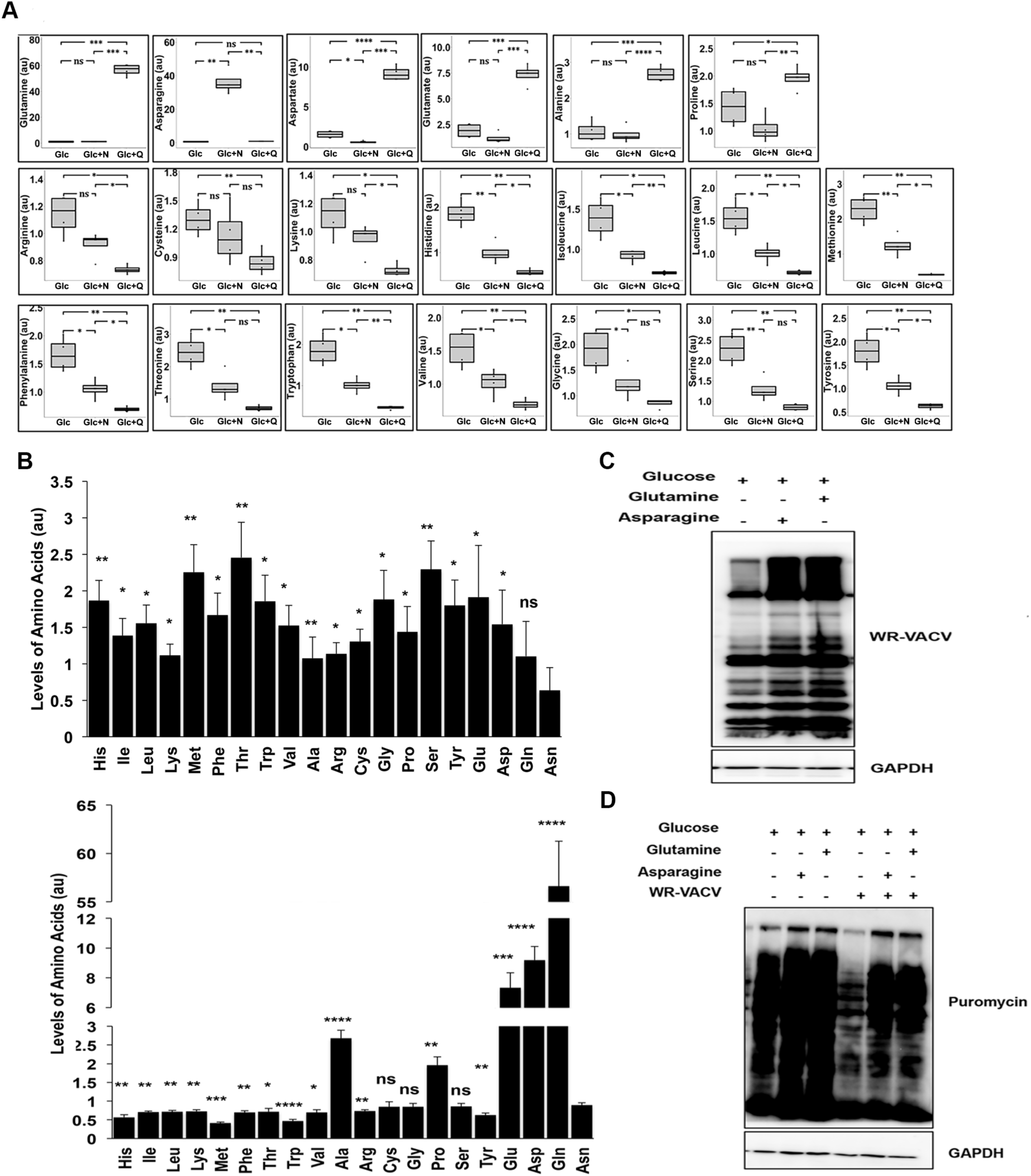
Asparagine rescues VACV protein synthesis from glutamine depletion. **(A)** Adding asparagine decreases accumulation of most amino acids, in glutamine-depleted conditions. Relative Levels of amino acids in HFFs infected with VACV, at an MOI of 3 for 8 h, in media as described in Fig 2A, as determined by a global metabolic profiling. **(B)** Upon glutamine depletion, asparagine is the least abundant amino acid in VACV infected cells. Upon glutamine repletion, asparagine is one of the most abundant amino acids. The levels of amino acids in HFFs infected with an MOI of 3 for 8 h in media with glucose (up) and glucose plus glutamine (bottom) was determined by global metabolic profiling. Statistical significance is shown by comparing to asparagine level. **(C)** Asparagine rescues VACV protein synthesis from glutamine depletion. Western blotting analysis was carried out in HFFs infected with VACV at an MOI of 2 for 24 h in indicated medium. GAPDH was used as a loading control. **(D)** Rescue of nascent protein synthesis by asparagine under glutamine-depleted conditions. HFFs infected with VACV, at an MOI of 2, or mock-infected for 16 h in the indicated medium. Cells were treated with 10 μg/mL puromycin for 10 min before collection, followed by Western blotting analysis using anti-puromycin and anti-GAPDH antibodies (the latter as a loading control). Error bars represent the standard deviation between four biological replicates. ns=p > 0.05, *= p ≤ 0.05, **= p ≤0.01, ***= p ≤0.001 and ****= p ≤0.0001.

For VACV-infected HFFs cultured in glutamine-deficient medium, adding asparagine did not increase the glutamine and glutamate concentration either (**Fig. 3A**). Moreover, an inhibitor of *de novo* glutamine synthesis, L-MSO, only minimally reduced VACV replication (1.4-fold) when glutamine-deficient medium was supplemented with asparagine (**Fig. 2F**), showing VACV replication is not rescued because asparagine increase glutamine or glutamate to feed the TCA cycle.

### Asparagine rescues VACV protein synthesis from glutamine depletion

Notably, our global metabolic profiling data showed most amino acids (14 out of 20) accumulated in cultures with glucose only, compared to cultures containing glutamine and glucose (**Fig. 3A**). With glucose in the medium only, amino acids whose biosynthesis is closely tied to glutamine concentration (i.e., alanine, proline, aspartate, glutamate and asparagine) ^30,31^ had a lower or similar concentration (**Fig. 3A**). Among the five amino acids, asparagine is the only amino acid that exclusively requires glutamine for its biosynthesis is also the only amino acid that fully rescues VACV replication from glutamine depletion. Remarkably, adding asparagine significantly decreased accumulation of most amino acids in glutamine-depleted conditions (**Fig. 3A**). Moreover, asparagine concentration was significantly lower than other amino acids except glutamine in glucose only condition, while its level was significantly higher than or similar to most other amino acids when glutamine is present (**Fig. 3B**).

These results prompted us to hypothesize that asparagine availability is a critical limiting factor for efficient protein synthesis in VACV-infected cells. This implies that, without glutamine, the rate of protein synthesis is actually suppressed by a low asparagine supply that cannot support the acute demand for nascent protein synthesis during the brief window of VACV replication. This hypothesis is supported by the result that VACV protein synthesis was much lower in cells cultured with glucose only, as compared viral protein levels with glutamine or asparagine (**Fig. 3C**). Consistently, the rate of nascent protein synthesis was also much lower in VACV-infected HFFs in medium containing glucose only, whereas in uninfected HFFs grown without glutamine, nascent, cellular protein synthesis was only mildly affected (**Fig. 3D**).

Phosphorylation of eIF2α is a key feature of global protein synthesis suppression in many biological process ^32^. However, we only observed a very mild increase of eIF2α phosphorylation in glucose only medium during VACV infection (**Fig. S2**). GCN2 is a metabolic-stress-sensing protein kinase that senses amino acid availability, and phosphorylates eIF2α to suppress protein synthesis ^33,34^. GCN2 phosphorylation was not evident either in glucose only medium (Not shown). Phosphorylation of S6K1, an indicator of the activation of mammalian target of rapamycin (mTORC1) that is another major pathway responding amino acid change for protein synthesis ^35^, was also only slightly increased in medium containing asparagine or glutamine during VACV infection (**Fig. S2**). In uninfected HFFs, these pathways were also only slightly affected or not affected (**Fig. S2**). This observation is consistent to the results that the most of amino acid levels increased in medium containing glucose only (**Fig. 3A**). Collectively, these results suggest that these amino acid sensing pathways do not play an important role in VACV protein synthesis rescue by asparagine from glutamine deficiency.

### Asparagine rescues VACV post-replicative mRNA translation from glutamine-deficiency

VACV genes are expressed in a cascade fashion, making it unclear at which step asparagine rescued VACV replication from glutamine-deficiency. Upon entry, VACV early genes are immediately expressed, then DNA is replicated and intermediate gene and then late genes are expressed ^5^. To find which phase was affected, HFFs were infected with one of three reporter VACVs that encode a secreted Gaussia luciferase gene under viral early (vEGluc), intermediate (vIGluc), and late (vLGluc) promoters. Viral gene expression was measured by Gaussia luciferase activities in cell culture medium. In all three conditions, early gene expression was similar but VACV intermediate and late gene expression was significantly higher when in medium containing asparagine or glutamine (**Fig. 4A**). qRT-PCR profiled VACV early (C11R), intermediate (G8R) and late (F17R) gene mRNA levels under different nutrient conditions (**Fig. 4B-D**) ^36–38^. Not surprisingly, the levels of early viral mRNAs were not affected by nutrient conditions (**Fig. 4B**). VACV intermediate mRNA levels did not differ either, and asparagine or glutamine only mildly increased viral late mRNA levels (**Fig. 4C-D**). These results indicate that asparagine rescues VACV protein synthesis from glutamine-deficiency at the post-replicative (both intermediate and late) mRNA translation stage.

**Figure 4.**
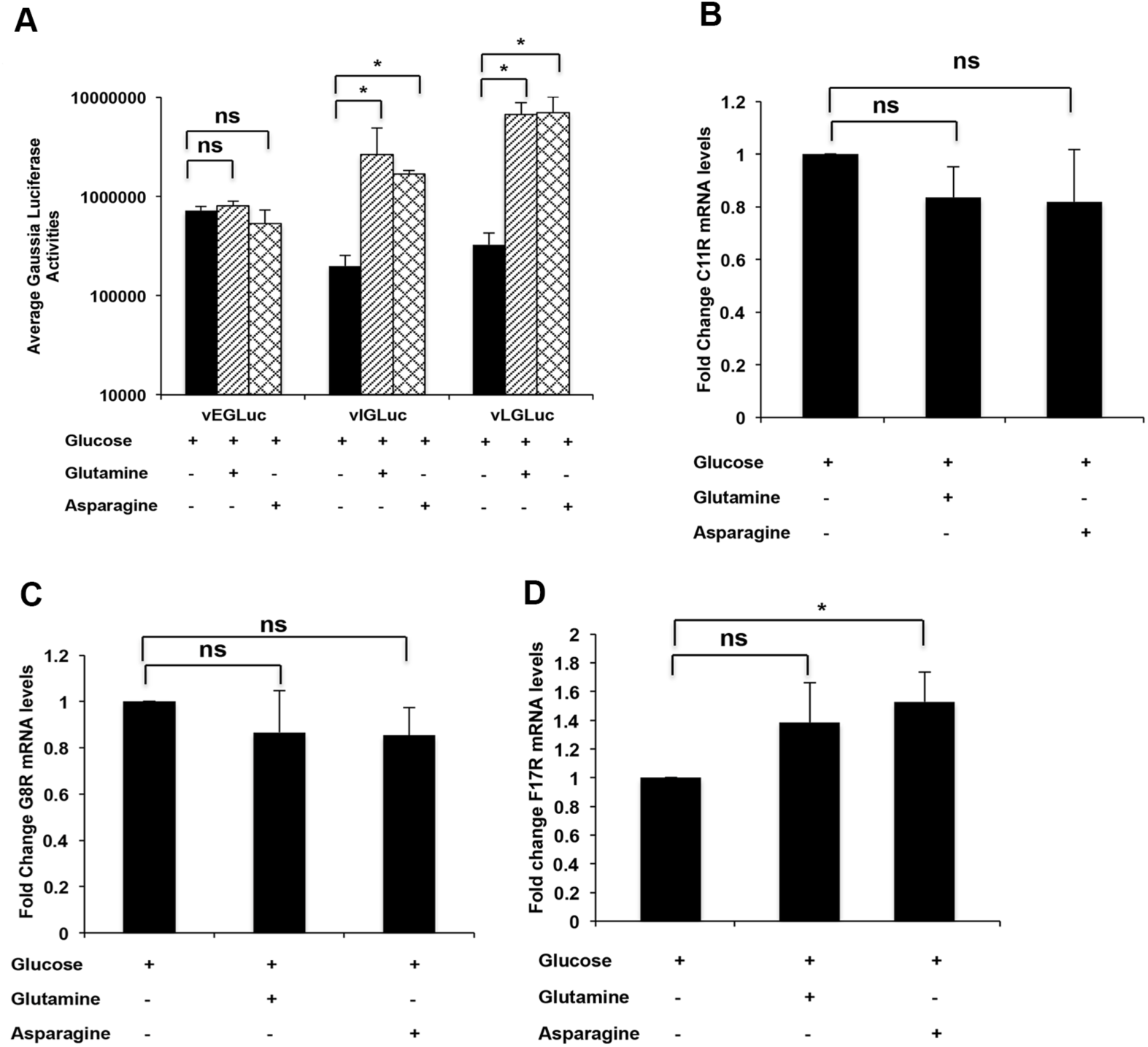
Asparagine rescues VACV post-replicative mRNA translation from glutamine depletion. **(A)** Efficient VACV intermediate and late gene expression, but not early gene expression, requires the presence of asparagine in glutamine-depleted medium. HFFs infected with VACV that expressed gaussia luciferase under early (vEGLuc; for 4 h), intermediate (vIGLuc; for 6 h), and late promoters (vLGLuc; for 8 h), respectively, in indicated medium, followed by gaussia luciferase activity measurement. **(B-D)** Addition of asparagine or glutamine in glucose only medium has no or mild effects on VACV early (C11R), intermediate (G8R) and late (F17R) mRNA levels. RNA was extracted from HFFs, infected with VACV at an MOI of 2, and qPCR was performed. Error bars represent the standard deviation between the at least three biological replicates. ns=p > 0.05, *= p ≤ 0.05.

Our metabolic profiling of cultures grown with glucose or asparagine supplementation showed that they had similar nucleoside concentrations (**Fig. S3A**). Accordingly, adding nucleosides to glutamine-depleted medium did not rescue VACV replication (**Fig. S3B**). These results support the conclusion that asparagine has little effect on VACV RNA production but significantly elevates post-replicative mRNA translation.

### ASNS knockdown impairs VACV replication

Standard cell culture medium lacks asparagine, and cells synthesize it *de novo* by ASNS, using glutamine as the amino group donor (**Fig. 5A**). To test whether asparagine biosynthesis affects VACV replication, ASNS protein expression was blocked with two specific siRNAs (**Fig. 5B**). ASNS knockdown significantly impaired VACV replication (**Fig. 5C, 5D**), but did not suppress cell proliferation (**Fig. S4**). In ASNS-siRNA–treated cells, VACV protein synthesis was down-regulated (**Fig. 5E**). This agrees with the result that siRNA-mediated interference of ASNS also decreased nascent protein synthesis in VACV-infected cells but not uninfected cells (**Fig. 5F**). Since these experiments were performed in the presence of glutamine, the results indicate a critical role of asparagine biosynthesis in VACV protein synthesis, and ultimately, viral replication.

**Figure 5.**
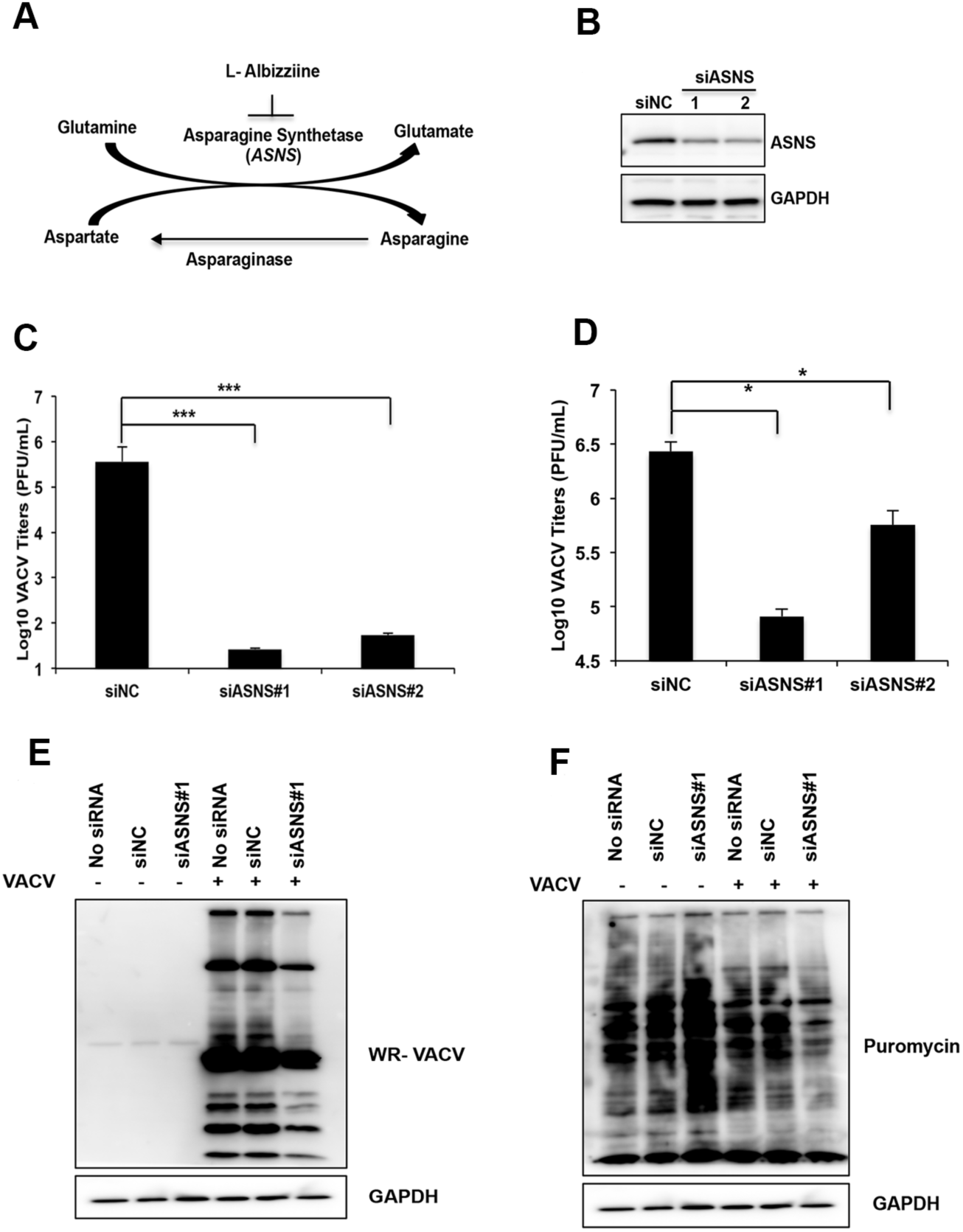
ASNS knockdown impairs VACV replication. **(A)** Schematic of asparagine metabolism. ASNS catalyzes the *de novo* biosynthesis of asparagine. Asparaginase catalyzes the conversion of asparagine to aspartate (inactive in mammalian cells). L-albizziine is a competitive inhibitor of ASNS. **(B)** ASNS siRNAs efficiently knock down ASNS protein level. HFFs were transfected with indicated siRNA for 72 h and the indicated proteins were detected using specific antibodies. **(C, D)** ASNS knockdown severely impairs VACV replication. HFFs were transfected with indicated siRNAs for 72 h and then infected with VACV at an MOI of 0.001 (C) or 2 (D). VACV titers were measured at 72 and 24 hpi, respectively, using a plaque assay. **(E)** ASNS knockdown does not inhibit nascent protein synthesis in uninfected cells but in infected cells. HFFs transfected with indicated siRNAs were infected with VACV at an MOI of 2 for 24 h. Cells were treated with 10 μg/mL puromycin for 10 min before the collection of cells for Western blotting analysis using indicated antibodies. **(F)** ASNS knockdown impairs VACV protein synthesis. HFFs transfected with the indicated siRNAs were infected with VACV, at an MOI of 2 for 24 h, and the proteins were analyzed by Western blotting analysis. Error bars represent the standard deviation between at least three biological replicates. *= p ≤ 0.05, ***= p ≤0.001.

### Chemically blocking asparagine metabolism inhibits VACV replication

Conversion of asparagine to aspartate can be catalyzed by asparaginase ^39^. To test how depleting asparagine affects VACV replication, asparaginase was added to culture medium containing glutamine. This significantly decreased VACV replication. Adding supplemental asparagine partially rescued VACV replication (**Fig. 6A**). Asparaginase treatment also decreased Gaussia luciferase activity in vLGluc-infected cells; and that also could be partially rescued with supplemental asparagine (**Fig. S5**). Although in medium containing asparagine, asparaginase treatment decreased VACV titers, in medium containing glucose only without asparagine and glutamine, asparaginase had no effect (**Fig. 6B**). Importantly, asparaginase treatment did not decrease cell viability (**Fig. 6C**).

**Figure 6.**
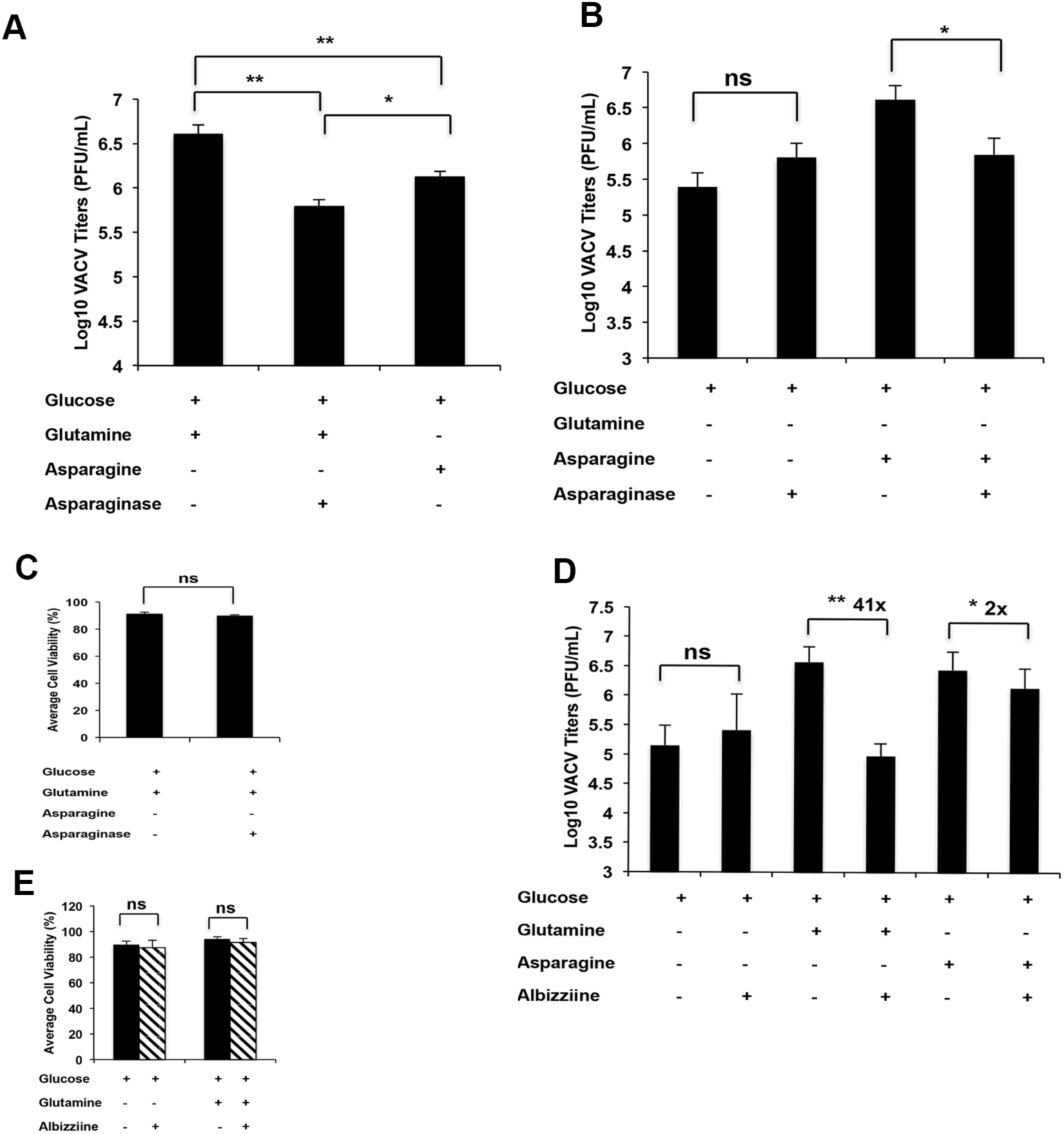
Interference of asparagine metabolism impairs VACV replication. **(A)** Asparaginase treatment reduces VACV replication. HFFs pre-treated with 10 U of asparaginase for 24 h and infected with VACV, at an MOI of 2 for 24 h, in indicated medium. Plaque assays measured VACV titers. **(B)** Asparaginase reduces VACV replication in medium containing asparagine but has no effect on medium containing glucose only without asparagine and glutamine. HFFs were treated with asparaginase and infected with VACV, at an MOI of 2 for 24 h, in indicated medium. Plaque assays measured VACV titers. **(C)** Asparaginase treatment does not reduce HFF cell viability. HFFs were treated with 10 U of asparaginase for 48 h before the cell viability were measured by Trypan blue exclusion assay. **(E)** Inhibition of ASNS by L-albizziine reduces VACV replication in glutamine containing medium. HFFs infected with VACV, at an MOI of 2 in, indicated medium in the presence or absence of 5 mM L-albizziine. VACV titers were measured at 24 hpi by a plaque assay. **(F)** Albizziine treatment does not reduce HFF cell viability. HFFs were cultured in indicated medium in the presence or absence of 10 mM L-albizziine for 24 h. Cell viability was measured by trypan-blue-exclusion assay. Error bars represent the standard deviation at least three biological replicates. ns=p > 0.05, *= p ≤ 0.05, ** = p ≤0.01. The numbers above the bars represent fold change.

To test whether chemical interference of ASNS impedes VACV replication, the ASNS competitive inhibitor, albizziine ^40^, was added to culture medium. In medium with glutamine, albizziine reduced VACV replication by 41-fold but had no obvious effect on VACV replication in cells grown with glucose only (**Fig. 6E**). In cells grown with asparagine plus glucose, albizziine reduced VACV titers by only two-fold (**Fig. 6E**). Furthermore, albizziine treatment decreased Gaussia luciferase activity in vLGluc-infected HFFs grown with medium containing glutamine and glucose, but not glucose only (**Fig. S6**). Albizziine had no effect on cell viability (**Fig. 6F**). Overall, these findings again demonstrate how interfering with asparagine metabolism impedes VACV replication.

## Discussion

This study used a combination of nutritional manipulation, and genetic and chemical approaches to establish that asparagine is a critical, limiting amino acid that controls VACV replication. During VACV replication, a rapid and heavy demand for the building blocks of protein synthesis triggers suppression of mRNA translation. During VACV infection in glutamine-containing medium, glutamine not only feeds the TCA cycle, it also is a substrate for asparagine synthesis, which supports the synthetic pathways supporting VACV replication (**Fig. 7A**). Because *de novo* synthesis of asparagine uses glutamine as the amino-group donor, asparagine cannot be synthesized in the absence of glutamine—this renders asparagine a limiting metabolite that must be exogenously supplied (**Fig. 7B**). In the absence of glutamine, a carbon source like glucose is required to feed the TCA cycle (**Fig. 7B**; in mammalian cells, asparagine is unable to feed the TCA cycle by converting to aspartate). When the asparagine supply is blocked by chemical or genetic approaches, VACV replication is suppressed, supporting our idea that asparagine is an essential metabolite limiting VACV replication (**Fig. 7B**). Thus, glutamine provides both the functions of glucose and asparagine required during VACV infection.

**Fig 7.**
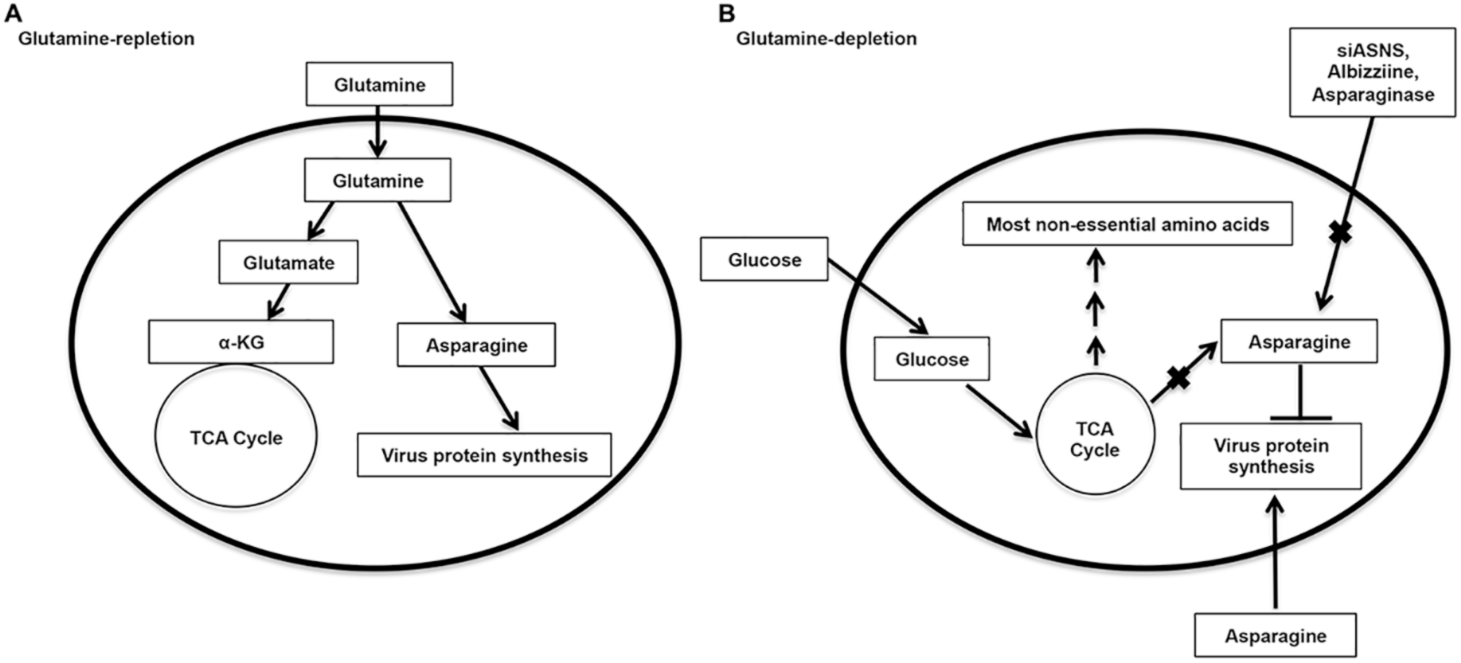
Proposed model for the role of asparagine in VACV replication. **(A)** In VACV infected cells under glutamine-replete conditions, glutamine supports the TCA cycle activities (glucose is dispensable) and also promote viral protein synthesis via asparagine *de novo* biosynthesis. **(B)** Under glutamine depletion conditions or other ways to interfere asparagine supply, VACV post-replicative protein synthesis is inhibited although glucose and/or glutamine are able to sustain the TCA cycle activities.

Glucose and glutamine are the two most common carbon sources for mammalian cells ^41,42^. Our findings support the idea that VACV’s preference of glutamine compared to glucose is accounted for by the requirement of glutamine during asparagine biosynthesis. Glutamine is sufficient for efficient VACV replication, even in the absence of glucose, suggesting glutamine can provide the function of glucose in VACV-infected cells. This observation led to the hypothesis that glutamine is the preferred carbon source during VACV infection. Yet, supplementation of α-KG and glutamate only partially rescued VACV replication from glutamine depletion; and, in the absence of glucose, glutaminase inhibition had a more severe effect on VACV replication (**Fig. 2C**). These results indicate that utilization of glutamine to feed the TCA cycle only partially accounts for its role during VACV replication and is not the cause of the glutamine preference during VACV replication.

Asparagine does not recapitulate glutamine’s effect on TCA cycle activity in VACV-infected cells, which might be explained by the lack of cytosolic asparaginase activity in mammalian cells ^28^. This idea is supported by our results that asparagine supplementation does not increase the aspartate level (**Fig. 3A**), and that aspartate in the absence of glutamine only weakly rescued VACV replication (**Fig. 2D**). In addition, asparagine needed glucose to rescue VACV replication from glutamine depletion (**Fig. 2E**). This is further supported by global metabolic-profiling data showing asparagine addition does not enhance TCA cycle activities in glutamine-depleted conditions (**Fig. 2A**). All of these facts support the idea that asparagine is not a carbon source during rescue of VACV replication from glutamine depletion.

Asparagine availability is critical for supporting VACV replication. Although glutamine can contribute to the biosynthesis of several non-essential amino acids via glutamate, asparagine biosynthesis exclusively requires glutamine ^23,24^. We showed that rescue of VACV replication from glutamine depletion is specific to asparagine (**Fig. 1D**). Inhibiting glutamine-dependent biosynthesis of asparagine (by genetic or chemical interference) severely impaired VACV replication, even if glutamine was present. Furthermore, asparaginase treatment, a well-established method of depleting asparagine ^43^, suppressed VACV replication. These results indicate that maintaining asparagine availability is essential for optimal VACV replication.

Asparagine mainly functions through its availability as a limiting nutrient for VACV post-replicative protein synthesis. The is supported by evidence showing an overall accumulation of most of the amino acids in glutamine depletion conditions, during VACV infection, suggesting they can be supplied by glucose metabolism. In contrast, asparagine does not accumulate and supplemental asparagine reduced the accumulation of amino acids in glutamine-depleted conditions. Interestingly, in uninfected HFFs, knockdown of ASNS or glutamine depletion has no or only mild effects on cell proliferation and protein synthesis. This is likely because the demand of nascent protein synthesis in uninfected HFFs (and during early stage of VACV gene expression) is lower. Because many viral infections demand rapid and robust protein synthesis to build up viral particles, asparagine might also be a limiting metabolite in the replication of other viruses.

Although asparagine is a protein building block, emerging evidence demonstrates that it plays a unique and specialized role in regulating various biological processes and disease development in mammalian cells ^30,44–46^. In mammalian cells, asparagine biosynthesis and metabolism are evolutionarily tailored so that its supply is limited and highly regulated. ASNS is the only enzyme to catalyze asparagine *de novo* synthesis. In glucose metabolism, asparagine is a non-essential amino acid to be synthesized at the very end of the TCA cycle and the synthesis is exclusively glutamine dependent (**Fig. 1A**). Unlike in flies and worms, it is not used to feed the TCA cycle by converting to aspartate in mammalian cells (asparaginase is inactive in mammalian cells) ^47,48^. These unusual features render it an attractive and significant target in studying disease development and treatment. To this end, asparagine has received increasing attention recently, especially for its essential role in cancer development. A new study indicates that asparagine controls breast cancer metastasis in an animal model ^27^. Asparagine is also important for cancer cell proliferation in multiple cancer cells, especially in the absence of glutamine, due to the requirement for various functions of asparagine ^23,24,28,49,50^. The key role of asparagine bioavailability in cancer development might, at least partly, account for its limited supply in mammalian cells. Asparagine metabolism is also critical in vessel formation ^50^. This, together with its importance for VACV replication suggests asparagine metabolism is a critical limiting factor in multiple biological processes and diseases, highlighting the importance to study metabolism regulation of individual amino acids to understand their roles in various life processes.

Asparagine metabolism can serve as an attractive target for novel anti-poxvirus strategy development. In fact, L-asparaginase has been used to treat various cancers, including acute lymphoblastic leukemia (ALL), acute myeloid leukemia (AML), and non-Hodgkin’s lymphoma for decades ^51–54^. In the future, it would be interesting to investigate how modulation of asparagine metabolism might affect poxvirus infection in an animal model; and its role in cancer cell proliferation and cancer development implicates asparagine metabolism as target for designing improved poxvirus-based cancer therapies.

## Materials and Methods

### Cells and viruses

Primary Human Foreskin Fibroblasts (HFFs; kindly provided by Dr. Nicholas Wallace, Kansas State University) were cultured in DMEM (Fisher Scientific) supplemented with 10% Fetal Bovine Serum (FBS, Peak Serum), 2 mM Glutamine (VWR), and 100 U/ml of Penicillin, 100 μg/ml Streptomycin (VWR). BS-C-1 cells (ATCC CCL-26) were grown in EMEM (Fisher Scientific) supplemented with 10% FBS, 2 mM Glutamine, and 100 U/ml of Penicillin, 100 μg/ml Streptomycin. All cells were incubated at 37°C in an incubator with 5% CO_2_. VACV Western Reserve (WR) strain (ATCC VR-1354) was used in this study. Amplification, purification, infection, and titration of VACV were carried out using methods described elsewhere ^55^. Recombinant VACVs encoding a Gaussia luciferase gene under an early (C11R, vEGluc), intermediate (G8R, vIGluc), or late promoter (F17R, vLGluc) were constructed by Dr. Jason Laliberte at National Institute of Allergy and Infectious Diseases (NIAID) and generously provided by Dr. Bernard Moss (NIAID). Recombinant VACV encoding a Green Fluorescence Protein (GFP) was described elsewhere ^56^.

### Antibodies and chemicals

L-Glutamate, L-aspartate, L-Serine, L-Proline, L-Alanine were purchased from VWR and used at indicated concentrations. L-Asparagine, Dimethyl 2-Oxoglutarate (Dimethyl α-ketoglutarate), L-Methionine Sulfoximine (L-MSO) and puromycin were purchased from Sigma-Aldrich and used at indicated concentrations. Asparaginase was purchased from Sigma-Aldrich. Dimethyl Sulfoxide (DMSO) and L-albizziine were purchased from Thermo Fisher Scientific. The EmbryoMax Nucleosides (100x) solution was purchased from EMD Millipore.

Anti-GAPDH was purchased from Abcam. Antibodies against GCN2, Phospho-p70 S6 Kinase (Thr389), p70S6K, Phospho-eIF2α (Ser51), total eIF2α were purchased from Cell Signaling Technology. ASNS antibody was purchased from Proteintech. Anti-puromycin antibody was purchased from Sigma-Aldrich. Antibodies against the whole VACV viral particle were kindly provided by Dr. Bernard Moss.

### Glutamine depletion and rescue

For glutamine depletion, DMEM without glucose, L-glutamine, sodium pyruvate, and phenol red (Fisher Scientific) was used. This medium also lacks L-asparagine. The medium was supplemented with 2% dialyzed FBS (Fisher Scientific), to thoroughly deplete small molecules and amino acids while still providing other essential factors for cell growth. For glutamine depletion and rescue experiments, 1 g/L glucose (Fisher Scientific), 2 mM glutamine, and 2 mM L-asparagine or other metabolites were added to the medium when necessary. The cells were washed with 1x PBS (VWR) prior to VACV infection.

### Global metabolic profiling

HFFs were grown in T-175 flasks. At about 95–100% confluency, they were washed with 1xPBS twice and then infected with VACV at the multiplicity of infection (MOI) of 3 in different media. After 8 hours post infection (hpi) the cells were harvested by scraping and the pellet was washed twice with ice-cold PBS. The pellet was then dissolved in the extraction solvent (methanol) and stored at −80 °C until shipment to Metabolon Inc. (Durham, North Carolina) for metabolic profiling. All the metabolic profiling experiments were performed with four biological replicates.

Proprietary analytical procedures were carried out to ensure highest quality data after minimizing the system artifacts, misassignments, and background noise among the samples. The raw reads were first normalized in terms of raw area counts and then each biochemical was rescaled to set the median equal to one. Then, missing values were imputed with the minimum. Values for each sample were normalized by Bradford protein concentration in each sample. Each biochemical was then rescaled to set the median equal to one and again missing values were imputed with the minimum. Three-way analysis of variation (ANOVA) with contrast tests was performed to calculate the fold change of metabolites.

### Cell viability assays

For the trypan-blue exclusion assay, cell viability was measured as described elsewhere ^57^. Briefly, after treatment, cells of each well (12-well plate) were treated with 300 μl of trypsin and resuspended with 500 μl of DMEM by pipetting. Twenty μl of cell suspension was gently mixed with 20 μl of 4% trypan blue (VWR). The numbers of living and dead cells were counted using a hemocytometer. MTT Cell Proliferation Assay (Cayman Chemicals) was performed as per the manufacturer’s instructions. Briefly, equal number of cells were seeded in a 96-well plate, allowing to grow overnight in a 37°C incubator, followed by necessary treatments and the absorbance measurement at 570 nm using a microplate reader.

### Gaussia luciferase assay

Cells were infected with recombinant VACV encoding a Gaussia luciferase gene for indicated time periods. The activities of Gaussia luciferase in culture medium were measured at indicated hpi using a Pierce Gaussia Luciferase Flash Assay Kit (Thermo Scientific) and a luminometer.

### Western blotting analysis and nascent protein synthesis analysis

The procedure was described elsewhere with minor modifications ^58^. For Western blotting analysis, cells were collected and lysed using RIPA lysis buffer (150 mM NaCl, 1% NP-40, 50 mM Tris-Cl, pH 8.0). Cell lysates were reduced by 100 mM DTT and denatured by sodium dodecyl sulfate–polyacrylamide gel electrophoresis (SDS–PAGE) loading buffer and boiling for 5 min before SDS–PAGE. After electrophoresis the proteins were transferred to a polyvinylidene difluoride membrane (Fisher Scientific). The membrane was then blocked in TBS-Tween (TBST) [50 mM Tris-HCl (pH 7.5), 200 mM NaCl, 0.05% Tween 20] containing 5% bovine serum albumin (BSA; VWR) for 1 h, incubated with primary antibody in the same TBST/BSA buffer for 1 h, washed with TBST three times for 10 min/each time, incubated with horseradish peroxidase-conjugated secondary antibody for 1 h, washed three times with TBST, and developed with chemiluminescent substrate (National Diagnostics). The whole procedure was carried out at room temperature. Antibodies were stripped from the membrane by Restore buffer (Thermo Fisher Scientific) for Western blotting analysis using another antibody.

To label the newly synthesized proteins, 10 μg/mL of Puromycin (Sigma Aldrich) was added to the cells 10 min prior to sample collection. The cells were then harvested for immunoblotting using anti-puromycin antibody.

### Real-Time PCR (RT-PCR)

Total RNA was extracted using TRIzol reagent (Ambion) followed by purification using Invitrogen PureLink RNA Mini Kit (Thermo Fisher Scientific). The RNA was used to synthesize cDNA using SuperScript III First-strand synthesis kit (Invitrogen) according to the manufacturer’s instructions using random hexamer primers. Relative mRNA levels were quantified by the CFX96 Real-time PCR Instrument (Bio-Rad) with All-in-One 2x qPCR Mix (GeneCopoeia) and primers specific for indicated genes. The qPCR program was started with initial denaturation step at 95°C for 3 min, followed by 40 cycles of denaturation at 95°C for 10 s, annealing and reading fluorescence at 52°C for 30 s, and extension at 72°C for 30 s. 18S rRNA was used as normalization factor for different samples.

### RNA interference

Specific and negative control siRNAs (siNC) were purchased from Integrated DNA technologies (IDT). HFFs were transfected at a concentration of 5 nm using Lipofectamine RNAiMAx (Fisher Scientific), according to manufacturer’s instructions. Knockdown efficiency was measured by Western blotting analysis of protein levels.

### Statistical Analysis

Unless otherwise stated, the data represented are the mean of at least three biological replicates. For the analyses of global metabolic profiling, four biological replicates were used for each treatment and the data was analyzed in an R Studio (version 1.1.442). A two-tailed paired *t-test* was used to evaluate significance in the difference between two means. Error bars represent the standard deviation between the experimental replicates. The following convention for symbols is used to indicate the statistical significance ns=p > 0.05, *= p ≤ 0.05, **= p ≤0.01, ***= p ≤0.001 and ****= p ≤0.0001.

## Acknowledgement

We thank Dr. Nicholas Wallace (Kansas State University) for providing HFFs. We thank Dr. Bernard Moss for providing various materials and reagents. The work was supported by grants from the National Institutes of Health to ZY (R21AI128406 and a subproject of P20GM113117). AP is also supported by Johnson Cancer Research Center of Kansas State University.

---

**Fig S1.**
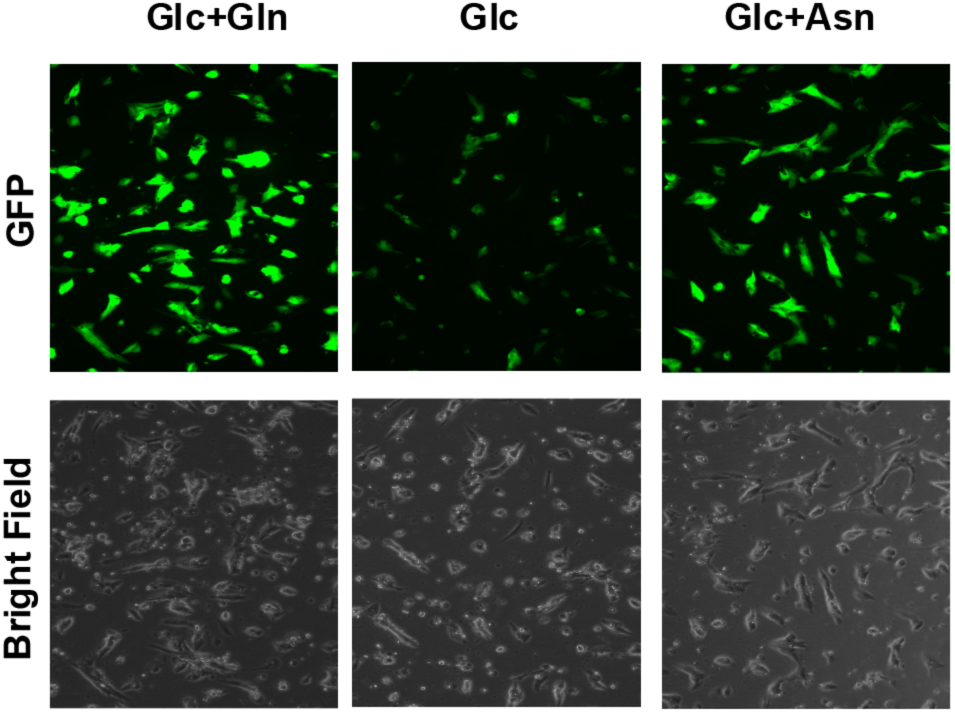
Asparagine rescues GFP expression from recombinant VACV in the absence of glutamine. HFFs infected with a recombinant VACV encoding GFP gene at an MOI of 2 in indicated medium. GFP expression was observed under a microscope at 24 hpi.

**Fig S2.**
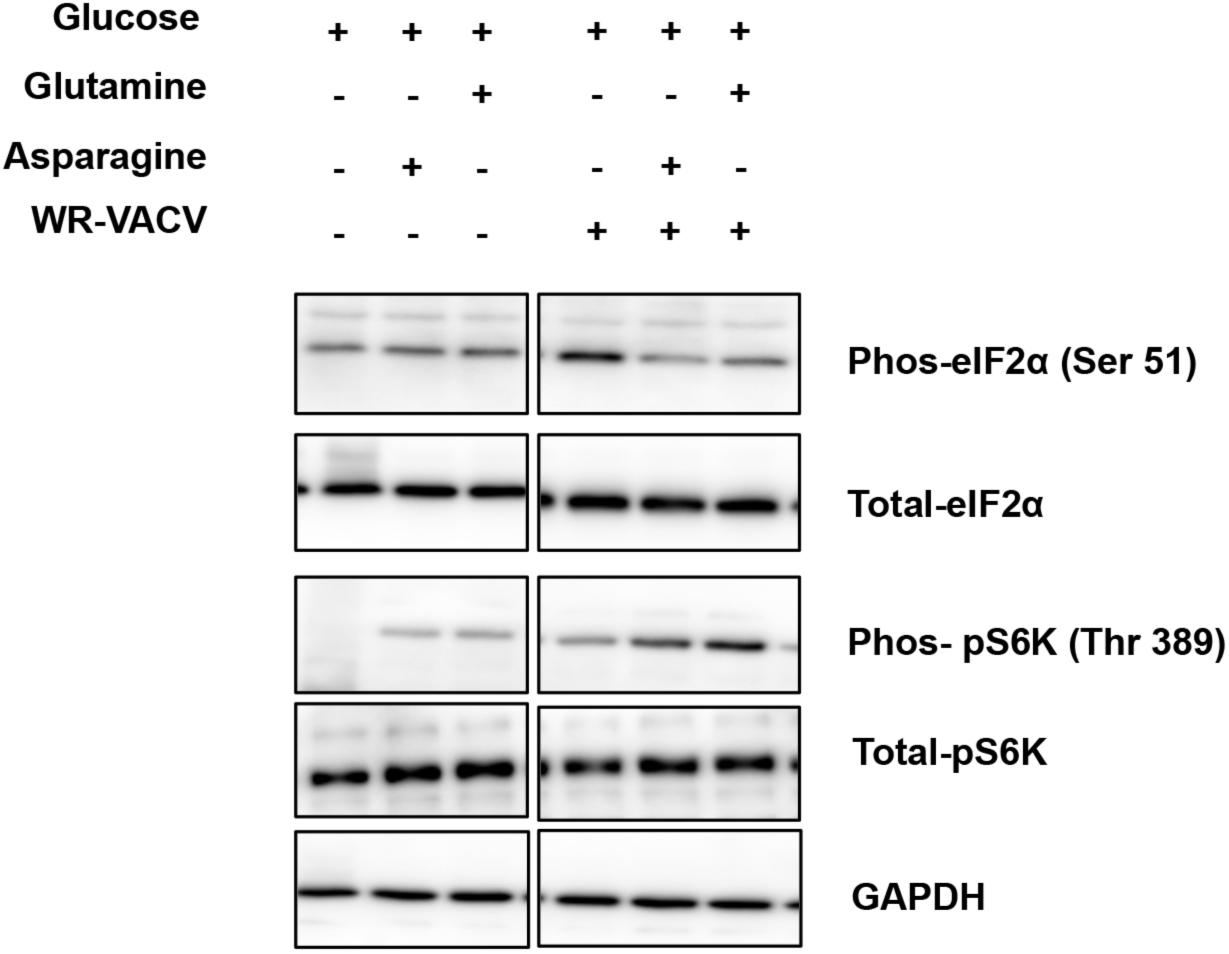
eIF2α and S6K1 phosphorylation is only mildly affected by different culture conditions containing glucose only, glucose plus glutamine, glucose plus asparagine during VACV infection. HFFs infected with VACV for 4 h, in the indicated media, and then levels of indicated proteins were determined by Western blotting analysis.

**Fig S3.**
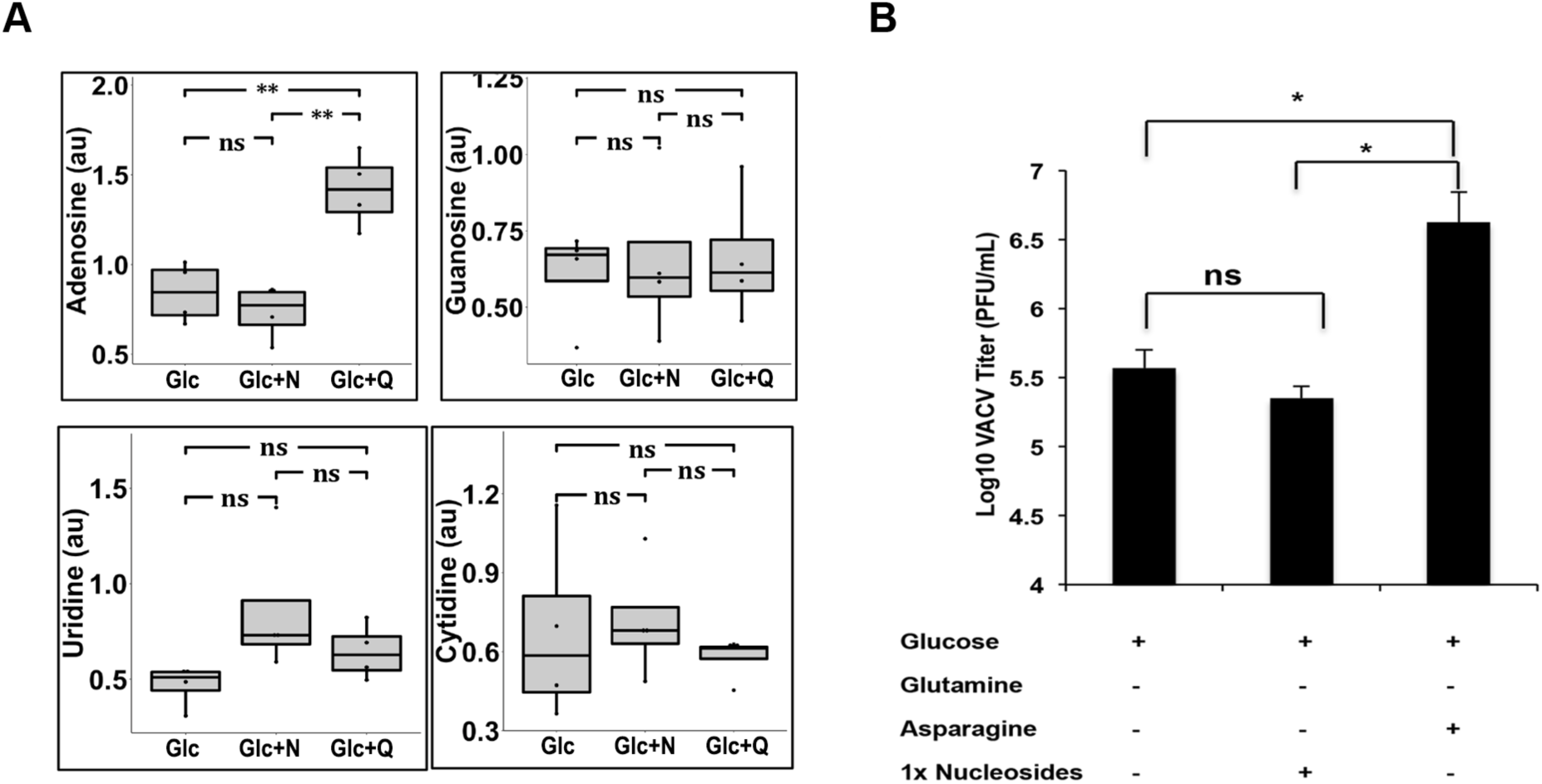
Asparagine-mediated rescue of VACV replication from glutamine depletion is not through nucleotide biosynthesis. **(A)** Asparagine does not increase levels of nucleosides in glutamine-depleted medium. Relative levels of nucleosides in HFFs infected with VACV, at an MOI of 3 for 8 h, in medium containing glucose (Glc), glucose plus asparagine (Glc+N) or glucose plus glutamine (Glc+Q), as determined by global metabolic profiling. **(B)** Addition of exogenous nucleosides in glutamine-depleted medium does not rescue VACV replication. HFFs infected with VACV at an MOI-2 in indicated medium for 24 h, followed by VACV titer measurement using a plaque assay. Error bars represent the standard deviation between at least three biological replicates. ns=p > 0.05, *= p ≤ 0.05, **= p ≤0.01.

**Fig S4.**
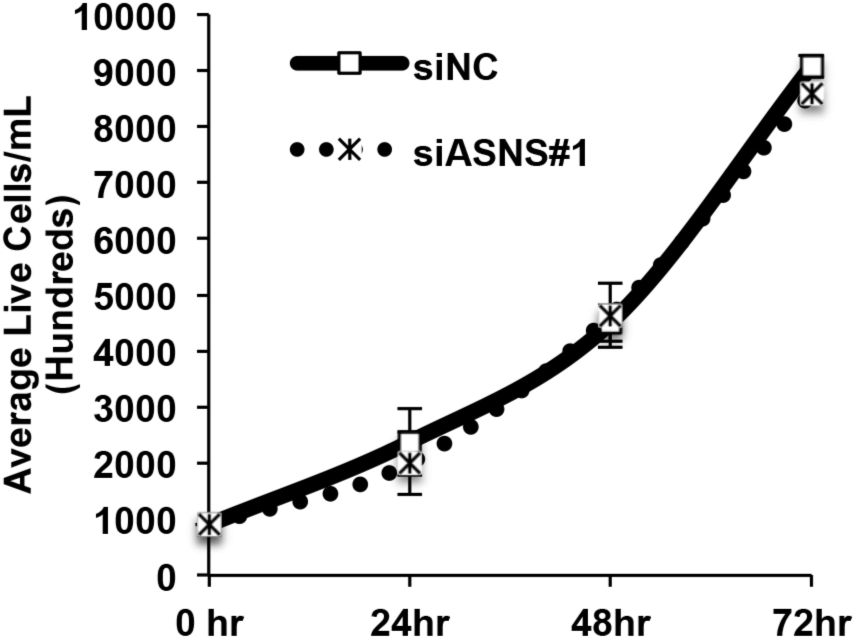
ASNS knockdown does not affect the proliferation of HFFs. HFFs treated with indicated siRNAs and numbers of live cells were counted using a hemocytometer for the indicated time period.

**Fig S5.**
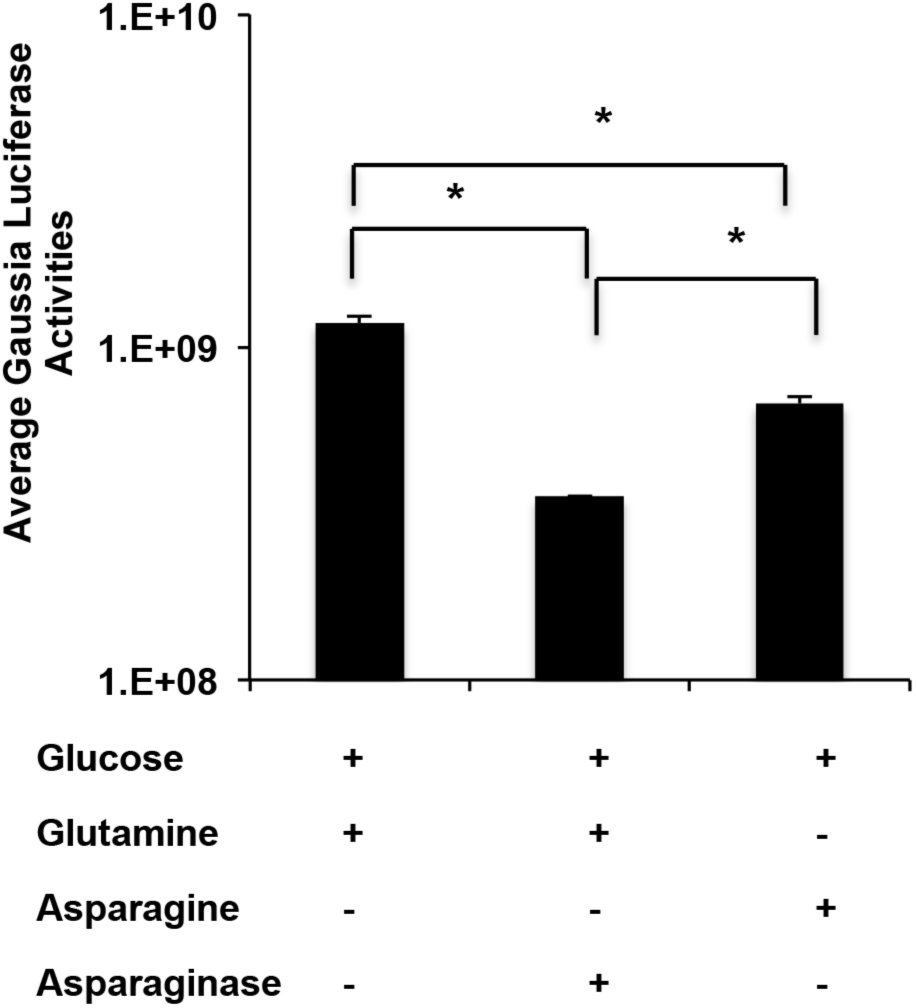
Asparaginase treatment reduces VACV gene expression. HFFs pretreated with 10 U of asparaginase for 24 h and infected with vLGLuc at an MOI of 2 for 16 h. Gaussia luciferase activities were measured. Error bars represent the standard deviation between at least three biological replicates. *= p ≤ 0.05.

**Fig S6.**
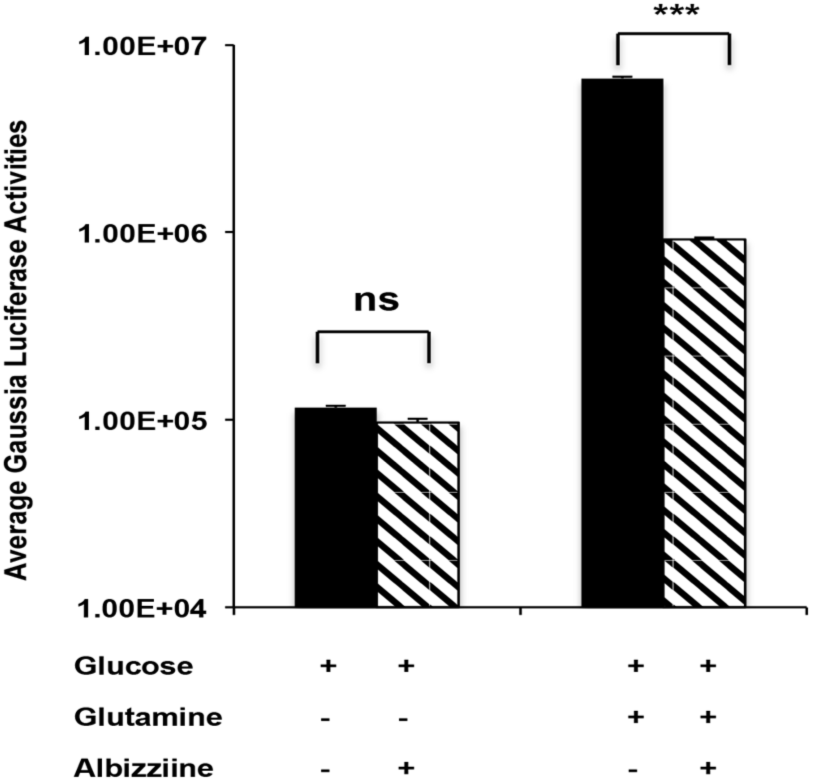
Albizziine reduces gaussia luciferase activity of recombinant VACV in glutamine-containing media. HFFs infected with MOI-2 of vLGLuc in indicated media in the presence or absence of 5 mM L-Albizziine. Gaussia luciferase activity was measured at 8 hpi. Error bars represent the standard deviation between at least three biological replicates. ns=p > 0.05, and ***= p ≤0.001.

